# The genomic basis of local adaptation to photoperiod across altitude in a self-fertilizing monkeyflower

**DOI:** 10.64898/2026.01.14.699501

**Authors:** Jill M. Love, Aditi Mahesh, Kathleen G. Ferris

## Abstract

Local adaptation along altitudinal gradients is well documented in many plant species, however the genetic basis of adaptive variation over these steep environmental clines remains poorly understood. Populations of *Mimulus laciniatus,* a self-fertilizing annual plant, experience highly differentiated seasonal environments throughout the Sierra Nevada, CA, where the length of the growing season and timing of favorable flowering conditions vary with altitude. These differences have driven divergence in critical photoperiod between low- and high-elevation *M. laciniatus*, an environmental cue that enables populations to initiate flowering at locally appropriate times. To investigate the genetic basis of local adaptation in this key ecological trait, we used a bulk-segregant quantitative trait locus (QTL) analysis approach. We crossed low- and high-elevation populations of *M. laciniatus* that differ in critical photoperiod to generate an F_2_ mapping population, phenotyping plants in a short-day common garden. Genomic differentiation (F_ST_ and *G*-statistic) between flowering and non-flowering pools identified 46 regions genome-wide associated with short-day flowering, including a strong peak on chromosome 8 overlapping *GA2ox3,* a candidate gene in the gibberellin pathway. Another gibberellin gene (*GA20ox2)* has been implicated in photoperiodic flowering in the close relative *Mimulus guttatus.* We found additional loci on chromosomes 2 and 11 that appear unique to *M. laciniatus*. Our findings suggest that local adaptation in reproductive timing may arise through a combination of shared genetic mechanisms and novel alleles in closely related Monkeyflowers, and that the genetic architecture underlying within-species adaptive divergence can be more complex than comparisons across species.

## Introduction

The timing of reproduction is a key life history trait known to be under strong selection across organisms in adapting to environmental heterogeneity. Reproductive synchrony influences what alleles are passed from one generation to the next and is critical during the process of local adaptation (Lande & Arnold, 1983). Theory predicts that adaptive shifts in reproductive timing can arise through diverse genetic architectures, ranging from single loci of large effect to highly polygenic networks of interacting genes, depending on the strength of selection, recombination, and demographic history (Yeaman & Whitlock, 2011). Rapid environmental change has increased recent interest in the genetic basis of reproductive timing, as temperature regimes, season length, and the reliability of environmental cues shift under climate change. This shift can generate mismatches between developmental cues and optimal reproductive windows, leading to detrimental fitness outcomes (Franks et al., 2014; Parmesan, 2006; Visser & Both, 2005). Together, these perspectives place reproductive timing at the intersection of life-history evolution, genetic architecture, and environmental change, making it an integrative lens for understanding how populations adapt to shifting seasonal regimes.

In annual plants – which must germinate, grow, flower, and set seed within the bounds of a single season – the timing of floral induction is critical as even small deviations in this schedule will affect whether reproduction occurs during the plant’s lifetime. Many temperate plants rely on day length (photoperiod) as a consistent seasonal cue, initiating flowering only once the number of daylight hours exceeds a genetically encoded critical photoperiod (Song et al., 2015). Below this threshold, floral development is suspended and individuals may fail to flower entirely, resulting in a complete loss of reproductive output. As a mismatch between seasonal cues and developmental thresholds can be disastrous for plant fitness, this trait has been the focus of genetic and developmental research in model systems such as *Arabidopsis thaliana* and crop species (Andrés & Coupland, 2012; Blázquez & Weigel, 2000; Cockram et al., 2007; Jung & Müller, 2009), but relatively little is still known about natural genetic variation in photoperiod in non-model plant species. Understanding the environmental conditions that favor different photoperiod thresholds in natural populations, and the genetic mechanisms that enable or constrain this variation, is therefore essential for explaining how populations adapt to heterogeneous seasonal conditions.

Photoperiodic regulation of flowering integrates external environmental cues with internal developmental and physiological states to ensure that reproduction occurs under favorable seasonal conditions. In plants, the perception of day length occurs primarily in leaves, where photoreceptors and circadian clock components interact to measure the duration of light and darkness and transmit this information to the shoot apical meristem to initiate floral development (Andrés & Coupland, 2012; Song et al., 2015). This process involves the coordinated regulation of multiple genetic pathways that converge on floral integrator genes, translating photoperiod signals into determinate developmental transitions. Importantly, photoperiodic flowering responses are not governed by light perception alone, but are modulated by interactions with a plant’s hormonal pathways, epigenetic regulation, and developmental thresholds that collectively determine when plants flower (Blázquez et al., 2003; Simpson & Dean, 2002; Song et al., 2013). Variation at any point along this regulatory cascade can shift the photoperiod threshold required for flowering, generating heritable differences in seasonal timing among individuals and populations. As a result, photoperiodic control represents a complex, multi-layered trait that provides numerous potential targets for natural selection to shape adaptive variation in flowering phenology across heterogeneous environments.

Montane plant species are excellent examples of organisms that have adapted to a steep environmental gradient in seasonal variation. The timing of snowmelt and the length of the growing season differ across steep elevation, creating strong spatiotemporal variation in the seasonal cues that trigger flowering. For annual plants in the Mediterranean climate of the Sierra Nevada (CA, USA), populations at low elevations experience lower snowpack, earlier snowmelt, and subsequently, earlier onset of seasonal drought. On the other hand, high-elevation populations contend with high snowpack and delayed melt leading to a later onset of seasonal drought, but simultaneously face early freezes and snowfall in autumn. As a result, plant populations throughout the Sierra Nevada should adapt to distinct critical photoperiods that initiate flowering at the appropriate time for their particular altitudinal environment (Keller & Körner, 2003). These locally adapted photoperiod thresholds help synchronize reproduction with the brief window in which conditions permit growth and pollination across specific elevations. However, as climate warming drives earlier snowmelt, soil warming, and onset of seasonal drought, the relationship between temperature cues and day length may decouple. Populations whose critical photoperiod do not shift accordingly may either germinate into a growing season that is too short for successful reproduction or lose synchrony within and among populations, reducing reproductive output, pollination success, and long-term persistence.

The Cutleaf Monkeyflower, *Mimulus laciniatus* (syn. *Erythranthe laciniata*) is an annual plant endemic to the Sierra Nevada, CA. It is a member of the *Mimulus guttatus* species complex, a closely related and largely interfertile group of wildflowers that is an emerging ecological genomic model system (Wu et al., 2008; Yuan, 2019). *Mimulus laciniatus* is known for its characteristic colonization of harsh granite outcrop habitats which have shallow rocky soils with little moisture retention causing them to experience rapid onset of seasonal drought (Ferris and Willis 2018). Populations of *M. laciniatus* grow along an elevational gradient from approximately 900 to 3270 meters, along which they experience differential seasonal day length conditions. Due to the timing of seasonal snowmelt, the peak period of growth varies with elevation; individuals at lower elevations (approximately 1220 m) germinate and grow in February and March during shorter photoperiods, while those at higher elevations (above 2300 m) germinate later leading to flowering in June and July (Dickman et al., 2019; Dong et al., 2025; Ferris & Willis, 2018; Tataru et al., 2024). Furthermore, despite its showy, bright-colored flowers, *M. laciniatus* is largely self-pollinating and highly inbred (Ferris et al., 2014; Sexton et al., 2011). Previous work by Love and Ferris (2024) demonstrated that populations of *M. laciniatus* display significant differentiation in critical photoperiod across the species’ altitudinal range and are widely phenotypically plastic in life-history related traits such as flowering time. A simple genetic basis for variation in critical photoperiod has been identified for close relatives *Mimulus guttatus* and *Mimulus nasutus* (Fishman et al., 2013; Friedman & Willis, 2013).

Mating system is a central determinant of how genetic variation is structured within and among populations, with important consequences for local adaptation. Self-fertilizing species typically exhibit reduced effective population sizes, elevated homozygosity, and lower levels of standing genetic variation within populations compared to outcrossing taxa (Charlesworth & Wright, 2001; Hamrick & Godt, 1996). As a result, theory has long predicted that self-fertilizing lineages may face constraints on adaptive potential, particularly across heterogeneous environments where selection favors locally tuned phenotypes. (Glémin & Ronfort, 2013; Wright, 2008). In reality, many highly self-fertilizing plant species like *M. laciniatus* occupy broad environmental ranges and show clear signatures of local adaptation (Siol et al., 2010). In such species, strong population structure, reduced gene flow, and rapid fixation of locally beneficial alleles may facilitate adaptive divergence despite limited within-population variation. The tension between expected genetic constraint and observed ecological specialization provides a dynamic framework for examining how self-fertilizing species evolve adaptive responses to steep environmental gradients.

This study investigates the genetic basis of local adaptation in critical photoperiod across elevation within a self-fertilizing annual plant species. Our primary objectives are to 1) evaluate variation and divergence in important life-history and morphological traits between *M. laciniatus* populations adapted to differential day length conditions (low and high elevation); and 2) determine the genetic architecture of divergence in critical photoperiod between high and low elevation *M. laciniatus* populations. We use a combination of common garden experiments, controlled crosses, and a bulk segregant analysis approach to quantitative trait locus (QTL) mapping (Ferris et al., 2015; Michelmore et al., 1991) to describe phenotypic variation and the genetic architecture of local adaptation in a key life history phenotype across an elevation gradient in the Cutleaf Monkeyflower.

## Materials and Methods

### Creation of the QTL Mapping Population

To identify the genetic basis of local adaptation in critical photoperiod across the altitudinal range of *M. laciniatus*, we first set out to create a genetic mapping population through controlled crossing. We chose parental lines from two populations of *M. laciniatus* that differed in their critical photoperiod. For this effort, we referred to a previous study exploring phenotypic plasticity in response to photoperiod variation in a broad sampling of *M. laciniatus* genotypes from across the species’ geographic range (Love & Ferris, 2024). We identified one short photoperiod sensitive (12 hr days) parental genotype from the low elevation Hetch Hetchy, California (HEH-9) population, which is found at ∼1220 meters. The other parental genotype only flowered under long days (15 hr) and was chosen from high elevation Huntington Lake, California (HUL-20), which is found at ∼2300 meters. The native HEH population typically flowers in March or April, under a shorter photoperiod, while the native HUL population typically flowers close to the summer solstice in June and July, experiencing a longer photoperiod during its brief growing season (Dickman et al., 2019; Ferris & Willis, 2018). To create our F_2_ hybrid mapping population we crossed low-elevation HEH-9 with high-elevation HUL-20 using hand pollination, resulting in F_1_ hybrids of which we self-fertilized one to create 504 F_2_s. HEH-9 was the maternal parent, and we performed no reciprocal parental crosses.

To create our experimental cross between HEH and HUL *M. laciniatus* populations we planted 30 individuals of each parental line in moist Fafard 4B soil within 3-inch by 3-inch pots. Pots were permanently flooded, left in a stratification chamber for 11 days at 4°C, and misted daily. Afterward, we transferred trays to a Conviron growth chamber which provided a 15-hour day length with a constant temperature of 21°C. The individuals remained in a permanently flooded bench until germination, after which point, they were flooded for 7 hours daily. We promptly thinned the plants to ensure that only one individual grew in each pot. To prevent self-pollen contamination and ensure a successful cross we emasculated low-elevation parental buds under a microscope by manually removing the four reproductive anthers and tagged them. Following high-elevation parental germination, we isolated anthers from randomly chosen individuals and manually brushed their pollen onto the stigma of an emasculated low-elevation parental individual. Once fruits formed, individuals were not watered to ensure that seeds were dry for collection. The F_1_ generation underwent the same grow-out protocol as the parental line. They were cold-stratified at 4°C, transferred to a 15-hour day length chamber, and permanently flooded until germination. After germination, they were flooded for 7 hours daily and allowed to self-fertilize to create F_2_ seeds.

### Measuring Phenotypic Variation

To measure the phenotypic response to photoperiod variation in our parental lines and F_2_ mapping population, we planted 30 low-elevation parental replicates (HEH-9), 30 high-elevation parental replicates (HUL-20), and 504 F_2_ individuals in short (12 hour) days. The same germination and growth conditions were used here as in the parental grow-out described above. As individuals were slow to flower in the 12-hour condition, the photoperiod was increased to 12.5 hours on their 60^th^ day in the growth chamber. For each individual, we collected phenotypic data including date of germination and first flower, plant height, and floral morphology related to *M. laciniatus’s* self-fertilizing mating system: corolla width and length, stamen length, and pistil length (Fig. S1). The experiment was concluded 109 days after the individuals were transferred to the growth chamber.

We analyzed phenotypic data using R v4.5.1 (R Core Team, 2025) and performed a chi-squared analysis (p < 0.05) to evaluate whether observed patterns in F_2_ flowering conform to the expected 3:1 Mendelian ratio of a single dominant locus. Twenty F_2_ individuals had only produced buds when the grow-out concluded and were removed from the analysis. We then performed a phenotypic correlation analysis in the R package *corrplot* (Wei & Simko, 2021) to evaluate the relationship between short-day phenotypic traits in the F_2_ population. In addition, we used t-tests to evaluate the differences in mean values of plant height, germination time, flowering time, and several floral traits (corolla length and width, smallest and longest stamen length, and pistil length) between the parental lines and F_2_ population. To account for multiple comparisons, p-values were adjusted using a false discovery rate (FDR) correction across all t-tests.

We calculated the broad-sense heritability (H^2^) of our life history and morphological traits using the equation V_P_ = V_G_ + V_E_, where V_P_ is the total phenotypic variance in a trait in the F_2_, V_G_ is the variance due to underlying genetic factors, and V_E_ is the variance due to environmental factors (Falconer & Mackay, 1996). We calculated V_E_ as the mean phenotypic variance of the parental inbred lines. We calculated V_P_ for each trait using the phenotypic variance of our F_2_ population (Fishman et al., 2002). When variance could be calculated for both parental lines, their values were averaged for V_E_. Otherwise, V_E_ was simply the variance of the low-elevation parent. For certain traits, the variance of the parental trait was higher due to the presence of an outlier data point (designated as falling outside the interquartile range), so the data point was removed to ensure a positive heritability value as suggested in Alvarez Prado et al. (2019) and Lourenço et al. (2020).

### Pooled Library Preparation and Sequencing for QTL Mapping

To perform our bulk segregant QTL analysis we collected tissue from each F_2_ individual, prioritizing bud and green leaf tissue for DNA extraction. Tissue was flash frozen and stored at - 80 °C. Following this, we performed DNA extractions using a standard CTAB-chloroform protocol (Fishman, 2020). We quantified the DNA concentration using an adapted PicoGreen (Invitrogen) protocol (J. Coughlan et al., 2020) on a FLX800 (BioTek) Microplate Reader. To identify critical photoperiod quantitative trait loci (QTL) in *M. laciniatus*, we used a bulk-segregant analysis (BSA) approach which estimates single nucleotide polymorphism (SNP) allele frequencies between selected pools of individuals at the phenotypic extremes of a hybrid distribution (Ferris et al., 2015; Friedman & Willis, 2013; Magwene et al., 2011; Michelmore et al., 1991). We selected two pools from our F_2_ mapping population: 1) the short-day pool – all F_2_s that flowered under short days (n = 52) and 2) the long-day pool – a random sub-sample of non-flowering individuals (n = 140). We chose to subsample the 282 individuals that did not reach flowering under short days for the non-flowering pool because it is optimal to have a more similar number of individuals across pools (Hivert et al., 2018; Schlötterer et al., 2014). We added equal amounts of DNA from each sample to the two pools ensuring that each individual was equally represented within its respective pool. Pooled samples were sent to the Duke University Genome Sequencing and Analysis Core Resource facility and sequenced using Illumina NovaSeq X Plus. All individuals of each pool were sequenced at ∼1X coverage per individual (short-day pool - 52X vs. long-day pool - 140X).

### Read Mapping and Filtering

Following sequencing, pooled F_2_ samples were quality-filtered (Phred = Q20) and aligned to the *M. guttatus* MgTOL v5.0 reference genome (Joint Genome Institute, 2023) using *BWA-MEM* (Li & Durbin, 2009). *Samtools* (Danecek et al., 2021) was used to sort, index, and convert alignments to mpileup format. We used *PoPoolation2* (Kofler et al., 2011) to generate synchronized pileup (.sync) files from aligned reads. Prior to downstream analyses, we retained only SNPs that met minimum read count and coverage thresholds (minimum allele count = 6, minimum coverage = 10, maximum coverage = 200). Per-site allele frequencies were estimated from the .sync file and then aggregated into non-overlapping 1000bp windows, based on prior *Mimulus* work showing that linkage disequilibrium decays to background levels within ∼1 kb (Brandvain et al., 2014). Windowed allele frequency differentiation (ΔAF) values (non-flowering – flowering pools) were normalized by chromosome-specific read depth. To visualize broader genomic patterns and reduce stochastic noise inherent to pool-seq data, we applied a 0.7 Mb running median smoother to the windowed ΔAF estimates.

We quantified genomic differentiation using both F_ST_ and the *G*-statistic (*G*). F_ST_ between flowering and non-flowering pools was calculated from the synchronized pileup file using *PoPoolation2*, which estimates genetic differentiation using a heterozygosity-based (Nei, 1973) estimator designed specifically for pooled sequencing data (Kofler et al., 2011). We calculated F_ST_ in a sliding-window scale with 1000bp step per 5000bp window. To complement this, we calculated *G* using allele count information derived from SNAPE (Raineri et al., 2012) following code modified from Gould et al. (2017). *G* is a widely used metric of genomic differentiation in bulk-segregant and pooled F_2_ designs, as it provides a variance-standardized measure of allele frequency divergence that explicitly accounts for read-count sampling variance (Magwene et al., 2011). This is an important consideration when classical population-genetic assumptions do not apply, as in an F_2_ mapping population (de la Fuente Cantó & Vigouroux, 2022; Magwene et al., 2011). We again used a sliding-window approach with 1000bp step per 5000bp window to calculate *G*. We used genome-wide empirical *G* thresholds to identify candidate regions, defining significant intervals as those containing three or more consecutive windows exceeding the 0.1% *G* cutoff (*G* clusters). These *G* clusters provide a conservative set of genomic intervals enriched for loci contributing to flowering under short-day conditions.

We also cross-referenced *G* with F_ST_ peaks to strengthen candidate inference. We first tested for exact overlap between F_ST_ and *G* outlier windows using genomic interval intersection implemented in the *Bioconductor* R package *GenomicRanges* (Lawrence et al., 2013). To evaluate concordance at genomic-level spatial scales, we additionally quantified the proportion of F_ST_ outlier windows with a *G* outlier within ±1 kb, 2 kb, and 5 kb. Distances to the nearest G-statistic outlier were calculated using *GenomicRanges*. Finally, to assess broader-scale concordance, we tested for association between the number of F_ST_ and *G* outlier windows per chromosome using Spearman’s rank correlation.

### Critical Photoperiod Candidate Gene Annotation

Candidate regions within a padded 10kb of *G* clusters were mapped to annotated genes in the *M. guttatus* MgTOLv5.0 annotation (Goodstein et al., 2012). For each gene, putative *Arabidopsis thaliana* orthologs were identified through the MgTOLv5.0 annotation file, and functional information was retrieved from The Arabidopsis Information Resource (TAIR) (Rhee et al., 2003) and UniProt (UniProt Consortium, 2025). We screened candidate loci for gene homologs using PaperBLAST (Price & Arkin, 2017) and searched for Gene Ontology (Ashburner et al., 2000; Gene Ontology Consortium et al., 2023) terms associated with flower development, photoperiodism, and light signaling, as well as previously reported flowering-time regulators.

## Results

### Phenotypic ratios suggest a single dominant Mendelian locus

To assess whether flowering under short-day conditions exhibits simple Mendelian inheritance or reflects a more quantitative genetic basis, we examined segregation ratios in the F_2_ population and estimated broad-sense heritability (H²) for key developmental and reproductive traits. We observed that the ratio of flowering to non-flowering individuals under short days matched the expected 3:1 ratio of a Mendelian dominant locus (χ^2^= 3.5703, df = 1, p = 0.059); (Table S1). H^2^ was also calculated for each trait to determine the proportion of phenotypic variance attributable to genetic variation among individuals (Table 1). We found that several traits exhibited strong heritability: plant height (66.0% due to genetic variation), germination time (57.8%), and flowering time (58.3%). Floral morphology traits varied considerably in their degree of heritability, with longest stamen length (42.4%) being most heritable, followed by corolla length (21.7%), pistil length (14.2%), shortest stamen length (13.1%), and corolla width (10.0%). These results indicate that variation in flowering ability and associated morphological traits have a substantial genetic basis between low- and high-elevation *M. laciniatus* genotypes.

**Table 1.** Broad-sense heritability of all traits measured.

| Trait | Broad-Sense Heritability ($H^2$) |
| --- | --- |
| Plant height | 0.660<sup>†</sup> |
| Flowering time | 0.583<sup>‡</sup> |
| Germination time | 0.578 |
| Longest stamen length | 0.424<sup>†‡</sup> |
| Corolla length | 0.217<sup>†‡</sup> |
| Pistil length | 0.142<sup>†‡</sup> |
| Shortest stamen length | 0.131<sup>†‡</sup> |
| Corolla width | 0.100<sup>†‡</sup> |

<sup>†</sup>Outliers were removed from one or both parental data sets.
<sup>‡</sup>Evaluated using only the low-elevation parental variance.

### Phenological variation between parental lines and F_2_ population

We quantified the phenology and morphology of parental lines and F_2_ mapping population in a short-day common garden. We detected both phenological and morphological differences between our parental lines and the F_2_ mapping population (Table 2). In terms of phenology, we found that the high-elevation parent germinated faster while the low-elevation parent flowered more rapidly. The high-elevation parent germinated 0.40 days faster on average than the low-elevation genotype (t_31.059_ = 2.408, p = 0.044; Fig. 1A). On average, the F_2_ population germinated more quickly than the high-elevation parent by 0.60 days (t_30.37_ = 3.366, p = 0.005) and the low-elevation parent by 1.03 days (t_77.67_ = 24.13, p < 0.001) (Fig. 1A). For flowering time, the low-elevation parent flowered 18.32 days earlier than the average of the F_2_ population (t_74.537_ = -8.850, p < 0.001; Fig. 1B). Only a single individual of the high-elevation parental line flowered 104 days following its germination in short days, therefore we have no statistical inference for its flowering time.

**Figure 1.**
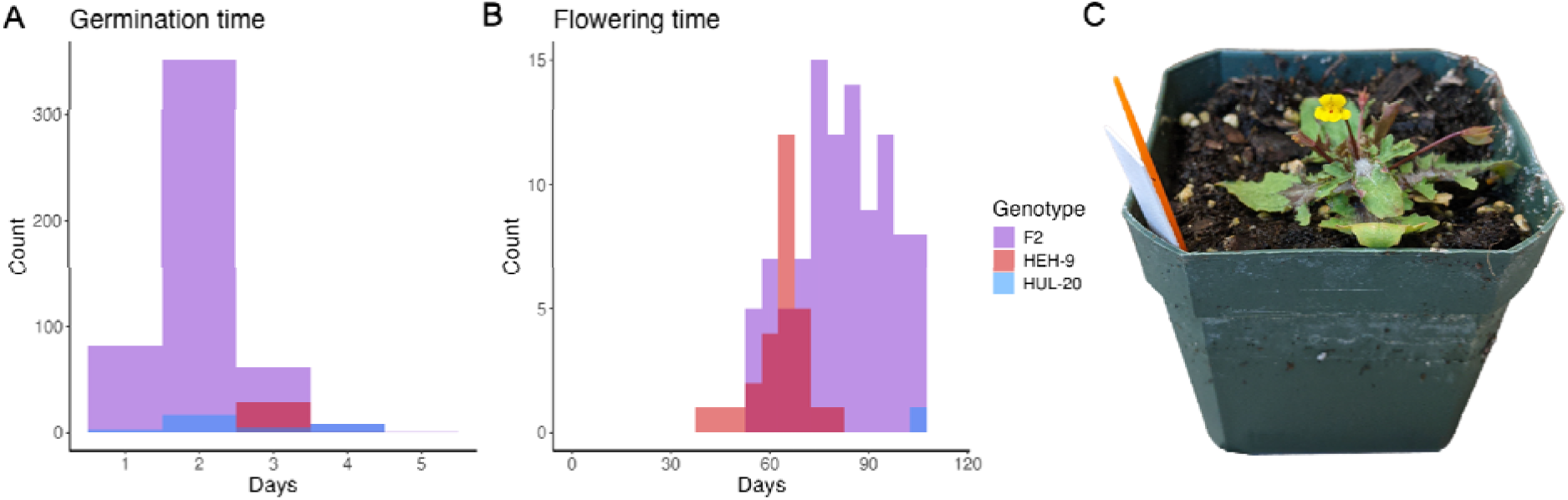
Distributions of phenological trait expression in the F_2_ mapping population (purple), low-elevation HEH-9 parental line (red), and high-elevation HUL-20 parental line (blue) under short days. **A)** Days elapsed between introduction to growth chamber and germination. **B)** Days elapsed between germination date and date of first flowering. **C)** Flowering F_2_ *M. laciniatus* in short days.

**Table 2.**
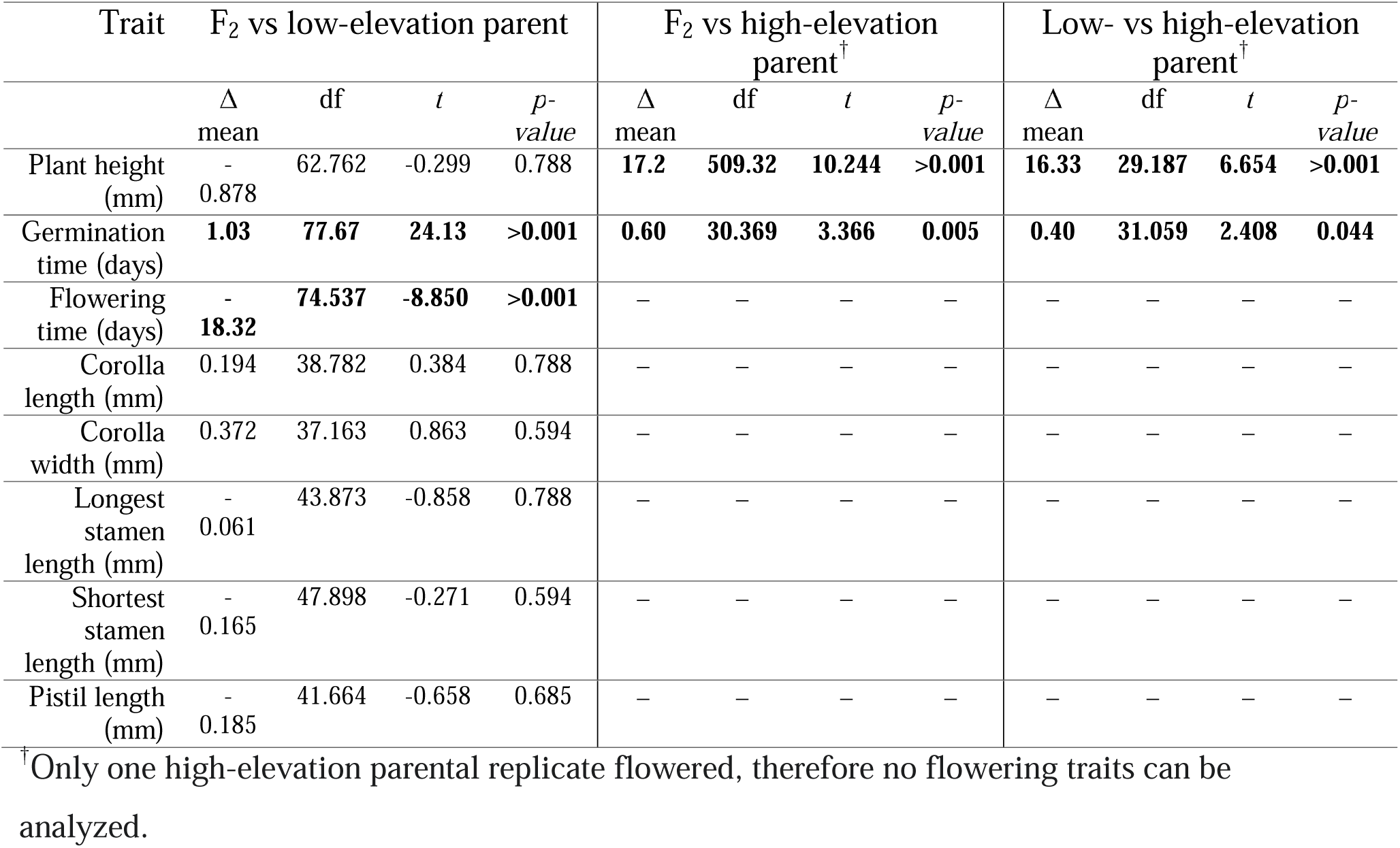
Differences in morphological and phenological traits between the low- and high-elevation parental lines and the F_2_ mapping population.

| Trait | F<sub>2</sub> vs low-elevation parent |  |  |  | F<sub>2</sub> vs high-elevation parent<sup>†</sup> |  |  |  | Low- vs high-elevation parent<sup>†</sup> |  |  |  |
| --- | --- | --- | --- | --- | --- | --- | --- | --- | --- | --- | --- | --- |
| | $\Delta$<br>mean | df | *t* | *p*-<br>*value* | $\Delta$<br>mean | df | *t* | *p*-<br>*value* | $\Delta$<br>mean | df | *t* | *p*-<br>*value* |
| Plant height (mm) | -<br>0.878 | 62.762 | -0.299 | 0.788 | **17.2** | **509.32** | **10.244** | **>0.001** | **16.33** | **29.187** | **6.654** | **>0.001** |
| Germination time (days) | **1.03** | **77.67** | **24.13** | **>0.001** | **0.60** | **30.369** | **3.366** | **0.005** | **0.40** | **31.059** | **2.408** | **0.044** |
| Flowering time (days) | -<br>**18.32** | **74.537** | **-8.850** | **>0.001** | - | - | - | - | - | - | - | - |
| Corolla length (mm) | 0.194 | 38.782 | 0.384 | 0.788 | - | - | - | - | - | - | - | - |
| Corolla width (mm) | 0.372 | 37.163 | 0.863 | 0.594 | - | - | - | - | - | - | - | - |
| Longest stamen length (mm) | -<br>0.061 | 43.873 | -0.858 | 0.788 | - | - | - | - | - | - | - | - |
| Shortest stamen length (mm) | -<br>0.165 | 47.898 | -0.271 | 0.594 | - | - | - | - | - | - | - | - |
| Pistil length (mm) | -<br>0.185 | 41.664 | -0.658 | 0.685 | - | - | - | - | - | - | - | - |

<sup>†</sup>Only one high-elevation parental replicate flowered, therefore no flowering traits can be analyzed.

Across all groups, we found that individuals that flowered were significantly taller than those that remained in their vegetative state (t_136.72_ = 10.395, p < 0.001) (Fig. S2). Following this, as all low-elevation individuals flowered, we found no significant difference between the average height of the F_2_ population and low-elevation parental line (t_62.76_ = -0.299, p = 0.788) (Fig. 2A). The high-elevation parental line was stunted in its growth, and on average, the high-elevation parent was shorter than the low-elevation parent by 16.33 mm (t_29.19_ = 6.754, p < 0.001) and the F_2_ population by 17.2 mm (t_509.32_ = 10.244, p-value < 0.001; Fig. 2A). This is an expected result, as flowering is a known trigger of plant height (Fabbro & Körner, 2004). The floral size traits (corolla length and width, longest and shortest stamen length, and pistil length) are all normal in distribution (Fig. 2B-F). There were no significant differences in average floral size between the F_2_ population and low-elevation parental line (p > 0.59 for all). Across all phenotypic traits in the F_2_ mapping population, we found strong positive correlation (*r* > 0.40) only between flowering time and plant height, with additional positive correlation among all floral size traits (Fig. S3).

**Figure 2.**
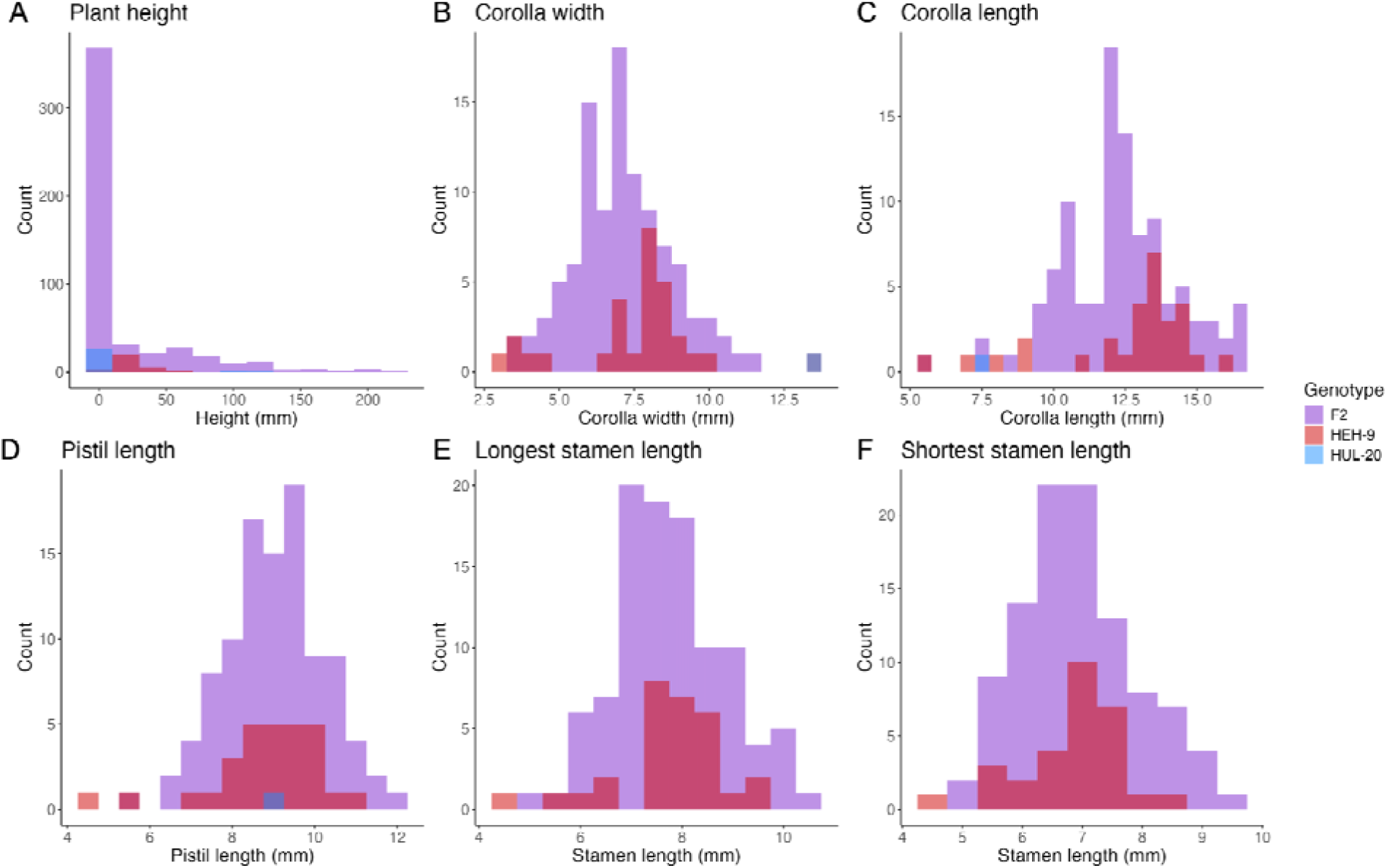
Distributions of morphological trait expression in the F_2_ mapping population (purple), low-elevation HEH-9 parental line (red), and high-elevation parental line (blue). Morphological traits measured were **A)** plant height, **B)** corolla width, **C)** corolla length, **D)** pistil length, **E)** longest stamen length, and **F)** shortest stamen length. Only one high-elevation parental individual (HUL-20) flowered during the experiment.

### Modest allele frequency differentiation between F_2_ sub-groups

To directly assess the magnitude and direction of allele frequency shifts associated with flowering under short-day conditions, we quantified genome-wide allele frequency differentiation (ΔAF) between flowering and non-flowering F_2_ groups. Genome-wide patterns of ΔAF revealed modest but structured divergence between the flowering and non-flowering F_2_ groups (Fig. 3A). Most of the genome exhibited near-zero mean ΔAF values, indicating limited allele frequency shifts between F_2_ groups. This is expected for any region unrelated to the trait we selected upon – critical photoperiod. Several regions did display localized deviations from zero, suggesting potential clusters of allelic differentiation related to flowering under short days. The strongest deviation occurred on chromosome 6, where normalized ΔAF magnitude exceeded the 99.9th percentile empirical threshold. This peak did not coincide with regions of allelic heterozygosity differentiation in F_ST_ (Fig. 3B) or *G*, suggesting that it does not reflect loci contributing to critical photoperiod divergence but rather stochastic variation, background linkage disequilibrium, or residual sampling noise (de la Fuente Cantó & Vigouroux, 2022; Magwene et al., 2011). Overall, the generally low genome-wide differentiation implies that only a small fraction of loci across the genome are associated with flowering under short-day conditions.

**Figure 3.**
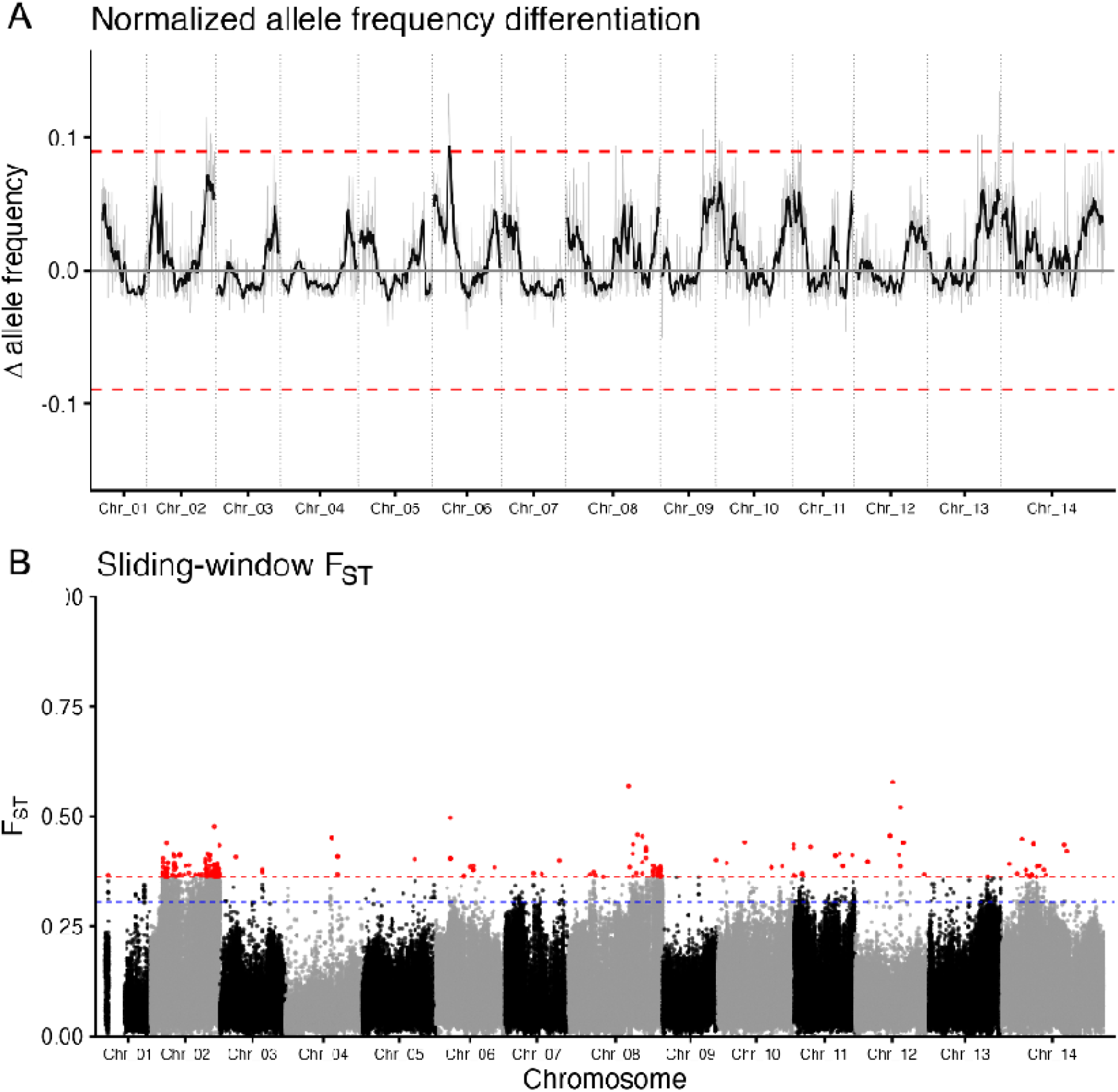
A) Genome-wide normalized allele frequency differentiation (ΔAF, in gray) between flowering and non-flowering F_2_ pools of *M. laciniatus*. The solid black line shows smoothed ΔAF (0.7 Mb median, ≥80 SNPs per window), and dashed red lines denote the 99.9th percentile empirical cutoff for extreme divergence. **B)** Genome-wide Manhattan plot of F_ST_ values comparing flowering and non-flowering F_2_ individuals. Each point represents a 1000bp genomic window, with alternating colors denoting chromosomes. The dashed blue line marks the empirical 99th percentile (top 1%), and the dashed red line marks the 99.9th percentile (top 0.1%) of F_ST_ values. Peaks above these thresholds are considered candidates for loci associated with critical photoperiod divergence.

### Candidate critical photoperiod genomic regions on chromosomes 2, 8, and 11

To identify genomic regions associated with critical photoperiod, we performed a bulk segregant analysis comparing allele heterozygosity differentiation (F_ST_) and *G*-statistic (*G*) values between flowering and non-flowering pools. In contrast to the expectations based on the phenotypic ratios in the F_2_ population, we found that critical photoperiod has a complex genetic basis with 246 candidate windows that exceeded 0.1% significance and 46 *G* clusters (Fig. 4).

**Figure 4.**
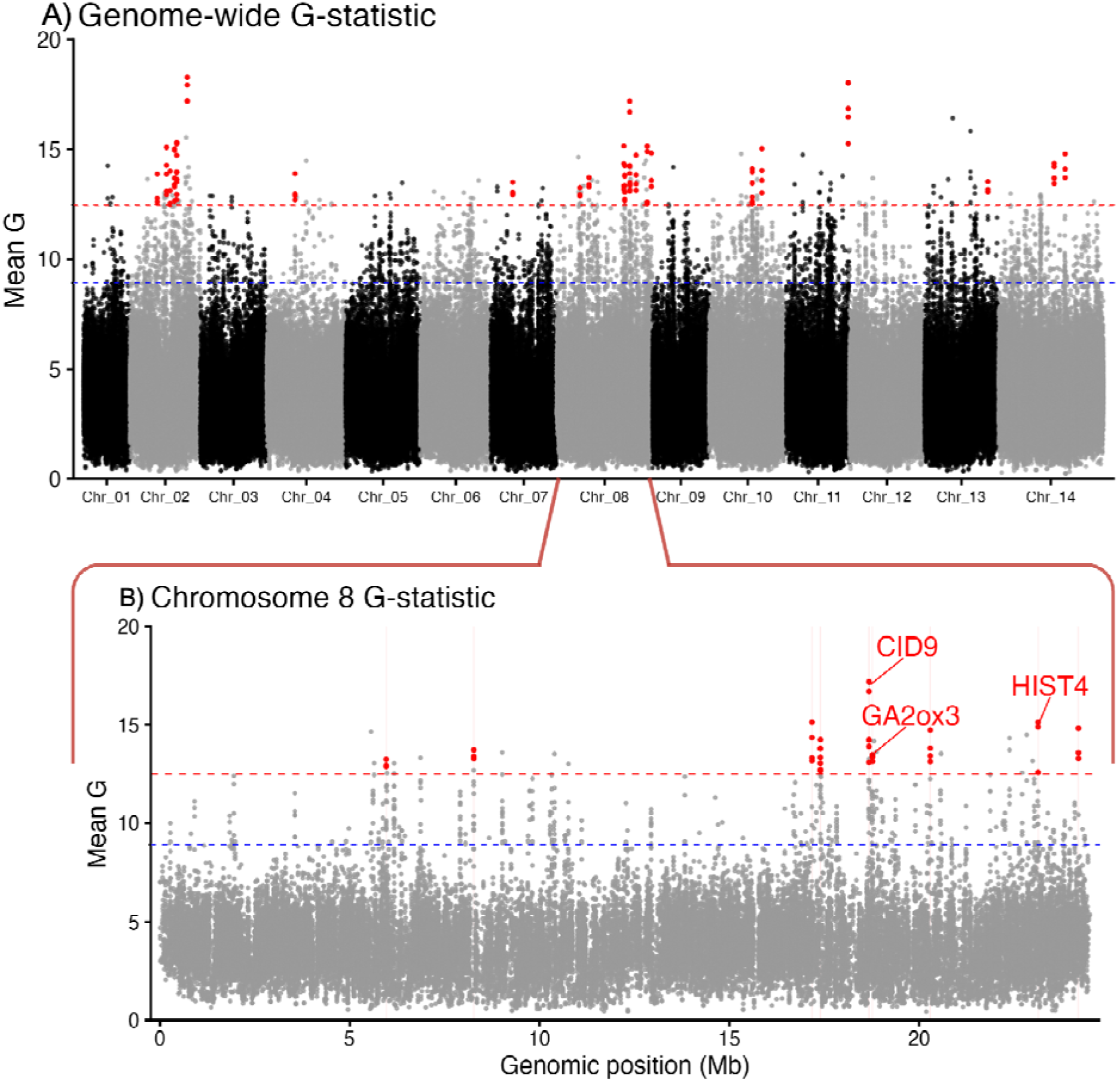
A) Genome-wide *G*-statistic (*G*) values comparing flowering and non-flowering F_2_ pools. Each point represents a 1000bp genomic window, with alternating colors denoting chromosomes. The dashed blue line marks the empirical 99th percentile (top 1%), and the dashed red line marks the 99.9th percentile (top 0.1%) of *G* values. 46 clustered peaks (red points) above the 99.9th percentile threshold were considered candidates for loci associated with critical photoperiod divergence. **B)** Chromosome 8 *G-*statistic with candidate genes. Candidate regions (±10kb from peaks) are highlighted in red bars.

We sought to determine overlap in outlier loci between our two metrics of differentiation. Despite visual co-localization of F_ST_ and *G* peaks in Manhattan plots, we observed no exact overlap between 1 kb outlier windows after aligning both statistics to a shared genomic grid. However, a small proportion of F_ST_ outliers occurred in close proximity to *G* outliers (1.6% within 1 kb; 5.3% within 5 kb); all of these proximity overlaps occurred on chromosomes 2 and 8. Together with a significant chromosome-level correlation in outlier density (Spearman’s ρ = 0.546, S = 206.44, p-value = 0.0433), these results indicate scale-dependent concordance: F_ST_ and *G* share peaks on the same chromosomes but typically on distinct, nearby areas rather than identical windows.

We chose to focus on *G* clusters in downstream gene annotation, as described in our methods, as *G* is a more statistically sound metric for pooled F_2_ genomic data. Notably, one of the highest *G* peaks lies on chromosome 8, similar to previous photoperiod QTL mapped in *M. guttatus* and *M. nasutus* (Fig. 4) (Fishman et al., 2013; Friedman & Willis, 2013). Other strong peaks on chromosomes 2 and 11 represent novel regions not previously associated with photoperiod sensitivity in *Mimulus*. These outlier regions represent QTLs contributing to variation in critical photoperiod. We found 46 genes located within a 10kb window around identified outlier regions (Table S1). After extracting gene functions from TAIR and UniProt, five genes stood out as candidate critical photoperiod regulators. Located on chromosome 2: delta-latroinsectotoxin-Lt1a protein (*Arabidopsis* ortholog AT3G08490; MgTOL.B1277); on chromosome 8: *GIBBERELLIN 2-OXIDASE 3* (*GA2ox3*; MgTOL.H2143), *CTC-INTERACTING DOMAIN 9 (CID9;* MgTOL.H2135), and *HISTONE 3 RELATED 4 (HIST4;* MgTOL.H2419); and on chromosome 11: galactose oxidase/kelch repeat superfamily protein (*Arabidopsis* ortholog AT1G80440; MgTOL.K1365). One additional candidate gene of note within the canonical *M. guttatus* chromosome 8 inversion interval (1.1–7.6 Mb) (Lowry & Willis, 2010) was identified (ribosomal L17 family protein; *Arabidopsis* ortholog AT5G64650; MgTOL.H0795).

## Discussion

Identifying the genetic architecture of local adaptation across environmental gradients is a major goal in evolutionary biology. Here we find that, contrary to our predictions based on previous *Mimulus* studies (Fishman et al., 2013; Friedman & Willis, 2013), divergence in critical photoperiod between low- and high-elevation *M. laciniatus* populations is a genetically complex trait. While the genomic architecture of critical photoperiod variation was more quantitative than expected, there were several large clusters of significant outlier windows on chromosomes 2, 8, and 11. In close relative *M. guttatus*, two large-effect QTLs on chromosome 8 explain most variation in critical photoperiod and vernalization between perennial and annual populations (Friedman and Willis 2013). While our results in *M. laciniatus* differ from these previous studies in the quantitative nature of the genetic architecture, we do identify strongly differentiated candidate genomic regions on chromosome 8 associated with flowering under short-day conditions (Gould et al., 2017; Kollar et al., 2025). Chromosome 8 is associated with divergence in many key life history traits across Mimulus species. However, as our chromosome 8 QTL do not precisely overlap with previously identified photoperiod QTL this indicates that within species divergence in critical photoperiod in *M. laciniatus* is genetically distinct from close relatives and that therefore there are many evolutionary paths to variation in photoperiod sensitivity at the molecular level. These findings contribute to a broader understanding of novel genetic components controlling photoperiodic flowering with implications for the evolution of seasonal timing, local adaptation, and mating system.

### Apparent Mendelian segregation masks a complex genomic architecture

We investigated the broad-sense heritability and phenotypic segregation ratios of key developmental and morphological traits that have diverged across an elevation gradient in the self-fertilizing *M. laciniatus*, including germination time, flowering time, and multiple floral size traits. Life-history traits exhibited high heritability (germination timing: 57.8%; flowering time: 58.3%), while estimates of heritability across morphological traits were more variable, ranging from 10% in corolla width to 66% in plant height, indicating a larger environmental influence on the expression of morphology under controlled conditions. At the phenotypic level, segregation of flowering versus non-flowering individuals under short-day conditions was consistent with a single dominant locus (Table S1). However, this apparent simplicity contrasts with the more complex genomic architecture revealed by our allele frequency and *G*-statistic analyses (Figure 4), which identified 46 differentiated regions between flowering pools across the genome. This discrepancy highlights how discrete phenotypic outcomes, such as flowering versus non-flowering, may mask underlying genetic complexity, particularly for traits governed by developmental thresholds and modifier loci. Such patterns are consistent with models in which a major-effect locus determines flowering status, while additional loci fine-tune flowering time and morphology among flowering individuals, a framework well supported in plant flowering-time regulation more broadly (Song et al., 2013, 2015).

We also quantified allele frequency differentiation (ΔAF) across F_2_ flowering groups to identify genomic regions associated with photoperiodic flowering. The modest magnitude and limited extent of ΔAF among flowering classes in the F_2_ mapping population highlight that, as expected, most of the *M. laciniatus* genome remains unassociated with photoperiod-induced flowering. The isolated ΔAF peak on chromosome 6, which did not coincide with a *G*-statistic outlier, may reflect recombination noise or linkage rather than selection on flowering under short days (Magwene et al., 2011). These results reinforce the idea that though flowering under short days is a genetically complex trait, adaptation to photoperiod across elevation in *M. laciniatus* involves a moderate number of loci rather than widespread genomic divergence, paralleling findings from related *Mimulus* species. More broadly, our results show support toward a growing body of literature that seasonal reproductive traits often exhibit genetically complex architectures. Across plant taxa – from herbaceous plants like *A. thaliana* to coniferous trees like lodgepole pine (*Pinus contorta*) – photoperiodic flowering has been shown to involve both major-effect loci and numerous modifiers, where local adaptation to seasonal cues arises from a multilocus gene framework rather than genome-wide divergence (Andrés & Coupland, 2012; Salomé et al., 2011; Yeaman et al., 2016). Similar patterns are present in animal response to seasonal environments as well (Bradshaw & Holzapfel, 2007). For example, diapause induction across dipteran, lepidopteran, and hymenopteran insects is frequently governed by a small number of key regulatory loci embedded in polygenic backgrounds that fine-tune developmental timing across latitudinal or climatic gradients (Ragland & Keep, 2017). Together, these comparisons suggest that adaptation to seasonal cues often occurs through a set of key loci operating within a complex genetic architecture, consistent with our findings in *M. laciniatus*.

### Candidate genes associated with hormonal regulation of flowering

The genetic basis for the transition from floral to vegetative growth has been explored in many plant species, such as *Populus trichocarpa* and *A. thaliana* (Böhlenius et al., 2006; Hsu et al., 2006; Koornneef et al., 1998). In *P. trichocarpa,* two genes, *PtFT1* and *PtFT2*, are light-dependent and serve as regulators for flowering time. Similarly, in *A. thaliana,* CONSTANS (*CO*) is a light-regulated locus that induces the transcription of genes in the flowering pathway (Suárez-López et al., 2001). In *M. guttatus*, critical photoperiod was found to vary extensively as a consequence of local adaptation, where two QTLs on chromosome 8 controlled critical photoperiod variation across populations (Friedman & Willis, 2013). Another factor in life history divergence and adaptation within *M. guttatus* is the chromosomal inversion on chromosome 8 (Gould et al., 2017; Lowry & Willis, 2010; Twyford & Friedman, 2015). While the inversion synteny is not currently known for *M. laciniatus*, within this historically defined inversion region (1.1 to 7.6 Mb) on chromosome 8 we identified one candidate gene, a ribosomal L17 family protein. This gene has been implicated in DNA and RNA methylation processes (Wan et al., 2015; Zhu et al., 2020), and not directly associated with flowering or photoperiod response (Rhee et al., 2003), suggesting it may play a more indirect or modulatory role, if any, in photoperiod-associated divergence.

Our genomic scan of short-day flowering versus non-flowering pools revealed strong allelic differentiation with the clearest candidate regions falling on chromosomes 2, 8, and 11, including three prominent regions on chromosome 8 (Fig. 4). This finding is interesting in the context of close relatives *M. guttatus* and *M. nasutus*, where two QTLs on chromosome 8 underlie floral morphology divergence between these close relatives (Fishman et al., 2002). Notably, we identified a candidate gene for critical photoperiod within the gibberellin (GA) hormone synthesis pathway, *GA2ox3* (Rieu et al., 2008). Another gene in the GA pathway also located on chromosome 8, *GA20ox2*, has previously been implicated in population divergence in critical photoperiod within the *M. guttatus* species complex (Gould et al., 2017; Kollar et al., 2025). Gibberellins are key regulators of flowering time and vegetative growth, integrating photoperiodic, thermal, and developmental signals to control the timing of reproductive transition (Mutasa-Göttgens & Hedden, 2009; Yamaguchi, 2008). Bioactive GA levels are determined by the balance between biosynthetic enzymes, such as GA 20-oxidases which promote the accumulation of GA precursors, and GA 2-oxidases which inactivate bioactive C19 gibberellins and thereby restrict GA signaling (Hedden & Thomas, 2012). *GA20ox2,* implicated in variation in critical photoperiod in *M. guttatus*, functions upstream in the pathway and modulates overall GA supply, with changes in its activity expected to broadly alter growth and flowering responses across tissues. In contrast, our candidate *GA2ox3* acts downstream by degrading bioactive GA, providing fine-scale spatial and temporal control over GA availability, and has been shown to respond to developmental regulators and photoperiod-associated signaling pathways (Rieu et al., 2008; Schomburg et al., 2003). Variation at *GA2ox3* in *M. laciniatus* may provide a mechanism for fine-scale, photoperiod-dependent regulation of bioactive gibberellin levels that govern the transition to flowering. We find that in *Mimulus*, photoperiod adaptation seems to repeatedly target the GA pathway, but different ecological contexts favor divergence at distinct regulatory nodes (Fig. 5). While *M. guttatus* has adapted differential flowering regimes across coastal versus inland ecosystems, *M. laciniatus* shows adaptation to flowering across an altitudinal gradient, with regulatory hormones that induce flowering in locally optimal conditions.

**Figure 5.**
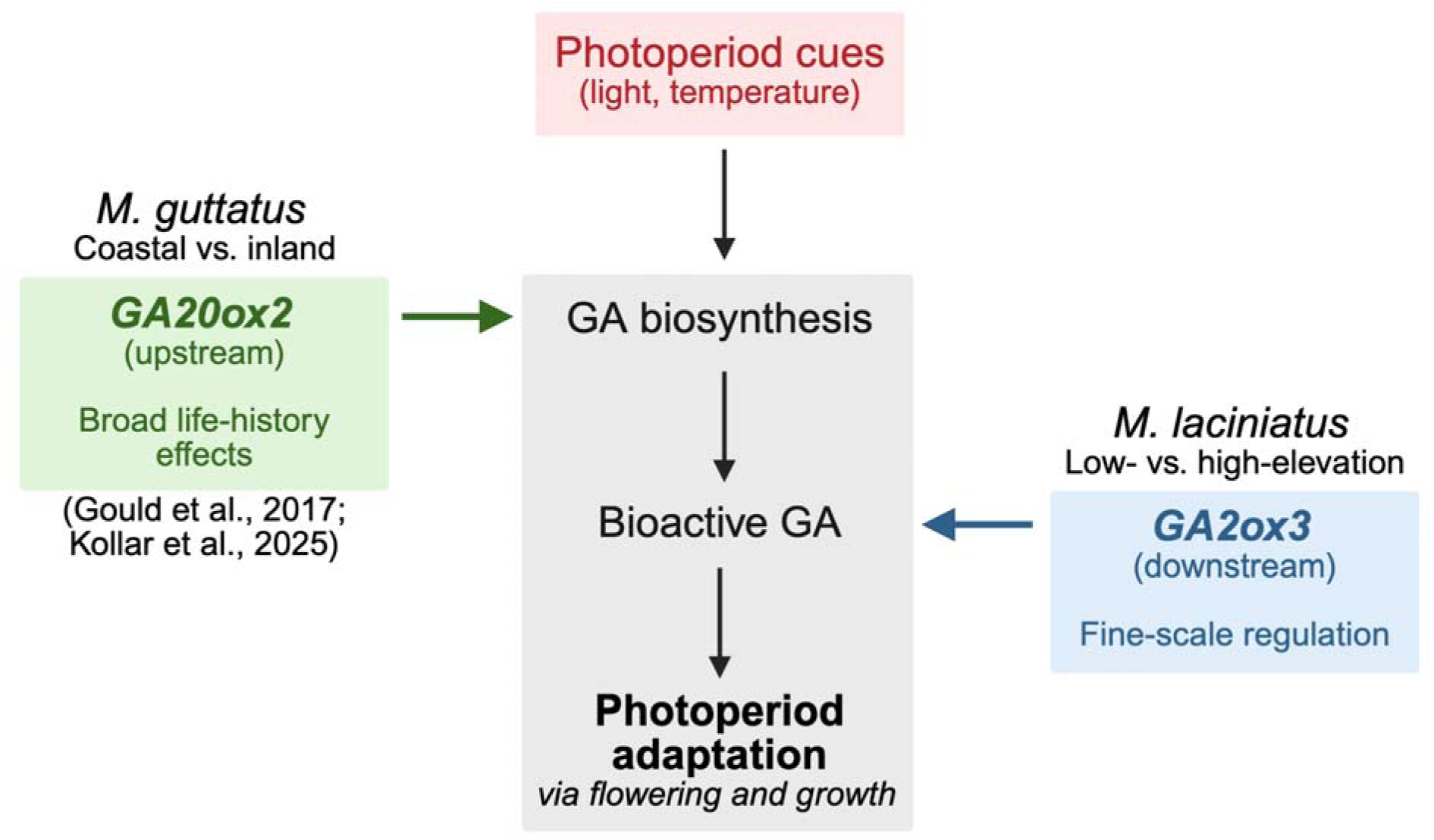
A conceptual diagram of the gibberellin (GA) hormone synthesis pathway utilized for photoperiod adaptation in both *M. guttatus* and *M. laciniatus*. Photoperiod cues trigger GA biosynthesis via GA 20-oxidase genes like *GA20ox2* in *M. guttatus*, while GA 2-oxidase genes like *GA2ox3* in *M. laciniatus* degrade bioactive GA for fine-scale regulation. Both approaches lead to photoperiod adaptation across divergent habitat types.

We also find several photoperiod candidate genes within our other QTL regions on chromosome 8: *HIST4* and *CID9* which have been found to influence light-regulated inflorescence development in *A. thaliana*. In *HIST4*, this is done primarily through chromatin-mediated control of flowering time (Whittaker & Dean, 2017; Zhao et al., 2021). The histone H3.3 family of genes, including *HIST4*, play an important role in transcriptional regulation by modulating chromatin accessibility, and have been shown to upregulate expression of the key flowering repressor gene *FLOWERING LOCUS C (FLC)* in *A. thaliana* (Zhao et al., 2021)*. FLC* is a central integrator of photoperiod, temperature, and stress cues, acting to delay flowering under non-inductive conditions (Michaels et al., 2003; Wilczek et al., 2009). Variation in chromatin-level regulation of *FLC* expression has been repeatedly implicated in local adaptation of flowering time (Michaels et al., 2003; Michaels & Amasino, 1999; Stinchcombe et al., 2004; Whittaker & Dean, 2017). In addition to its role in flowering repression, *HIST4* expression responds to environmental stress signals, including those associated with short photoperiod and seasonal transitions, suggesting that histone-mediated regulation may couple photoperiod sensing with broader stress-response pathways (Bao et al., 2016; Endler et al., 2015; Griffiths et al., 2024; Long et al., 2024; Mérai et al., 2019). Together, these findings support a model in which divergence at *HIST4* could influence photoperiod adaptation by altering chromatin states that regulate flowering repressors such as *FLC*, providing a mechanistically distinct but complementary route to photoperiodic divergence relative to hormone-based pathways. By contrast, *CID9* is a Class D CTD-interacting domain protein that remains poorly characterized functionally, currently defined primarily by conserved sequence features rather than direct experimental evidence (Bravo et al., 2005; López-Juárez et al., 2021). Although its precise role in transcriptional regulation is unresolved, the presence of CID9 within a QTL region presents it as an intriguing yet provisional candidate for contributing to photoperiod divergence in *M. laciniatus*, potentially through effects on transcriptional processing. Our results indicate that in *Mimulus* chromosome 8, which has been implicated in many aspects of life-history divergence in other members of the *M. guttatus* species complex (Coughlan & Willis, 2019), commonly harbors loci regulatoring photoperiodic flowering.

The remaining *M. laciniatus* QTL regions on chromosomes 2 and 11 were not previously associated with photoperiod traits in *Mimulus*, indicating novel lineage-specific loci contributing to flowering under short-day conditions at low-elevation. The genes delta-latroinsectotoxin-Lt1a protein (MgTOL.B1277) and galactose oxidase/kelch repeat superfamily protein (MgTOL.K1365) on chromosomes 2 and 11 respectively have functions that are less well understood; though for the galactose oxidase/kelch repeat superfamily protein, one study found differential gene expression related to sexual reproduction in the genus *Boechera* (Zühl et al., 2019). While the direct roles of these genes in photoperiod response remain unclear, the presence of differentiation in these regions alongside evidence of selective expression during *A. thaliana* flowering (Rhee et al., 2003) suggests that in *M. laciniatus*, multiple independent loci contribute additively or epistatically to flowering under restrictive short-day conditions. The presence of 43 smaller areas of allelic differentiation across the genome in addition to these three major peaks further suggests a more complex architecture than a single-locus model would imply. These may represent modifier loci. This complex architecture points to a nuanced genetic basis of photoperiod local adaptation within this highly self-fertilizing species.

### Genetic architecture of adaptation shaped by self-fertilization

Self-fertilization fundamentally alters the population structural and quantitative genetic contexts in which local adaptation evolves, influencing both the amount and distribution of genetic variation underlying adaptive traits. Classic population genetic theory predicts that self-fertilization reduces effective population size, increases linkage disequilibrium, and limits within-population standing genetic variation, potentially constraining adaptive responses to spatially variable selection (Charlesworth & Wright, 2001; Wright, 2008). From a quantitative genetic perspective, increased homozygosity in self-fertilizing lineages is expected to reduce additive genetic variance within populations while amplifying among-population divergence through drift and local fixation of alleles (Charlesworth & Charlesworth, 1995; Lande, 1976). These dynamics generate a genetic architecture of local adaptation characterized by strong population structure, reduced polymorphism within populations, and the potential for large-effect loci or tightly linked clusters of alleles to play a disproportionate role in phenotypic divergence (Glémin & Ronfort, 2013). Our results are consistent with these expectations: despite limited within-population variation (Awadalla & Ritland, 1997; Ferris et al., 2014), critical photoperiod divergence in *M. laciniatus* is associated with a genomic region on chromosome 8 plus multiple additional loci of smaller effect, suggesting that self-fertilizing does not preclude a complex genetic architecture. Instead, mating system may reshape how adaptive variation is partitioned across the genome. In this context, strong selection along steep elevation gradients may favor the rapid fixation of locally beneficial alleles and the accumulation of modifier loci, enabling fine-tuned phenological adaptation even in a highly self-fertilizing species (Glémin, 2012; Kawecki & Ebert, 2004).

## Conclusion

Our findings provide evidence for a genetically complex basis of critical photoperiod divergence across elevation in *M. laciniatus*, but with a strong *G*-statistic peak corresponding to a GA gene on chromosome 8 consistent with previous work in *M. guttatus* and *M. nasutus* (Fishman et al., 2013; Friedman & Willis, 2013; Kollar et al., 2025). These findings illustrate how phenotypic patterns of simple inheritance can coexist with underlying genomic complexity, and highlight the power of integrative approaches for revealing the full spectrum of genetic contributions to adaptive traits (Barghi et al., 2020). The combination of the gibberellin pathway being repeatedly co-opted to modify flowering across *Mimulus* species alongside novel genomic regions suggests that *M. laciniatus* populations balance robustness with flexibility in their flowering responses – an architecture well-suited to life along steep elevational gradients where the onset of favorable conditions can vary dramatically. As climate change reshapes the timing of snowmelt and seasonal cues, such distributed genetic control may beneficially influence the potential for populations to track shifting environments (Chevin et al., 2010). Our findings therefore highlight not only the genomic basis of photoperiod divergence, but also its broader role in mediating local adaptation and future persistence under heterogeneous seasonal regimes (Gomulkiewicz & Holt, 1995).

## Acknowledgements

Thank you to Natalie Gonzales for assistance with lab work, and Diana Tataru, Caroline Dong, and Jessica Scales for assistance with genomic prep. We thank the Duke University School of Medicine for the use of the Sequencing and Genomic Technologies Shared Resource, which provided their genomic sequencing service. This work was supported by the National Institute of General Medical Sciences of the National Institute of Health (NIH) under Award Number R35GM138224. The content is solely the responsibility of the authors and does not necessarily represent the official views of the NIH.

We acknowledge and pay tribute to the original inhabitants of the land Tulane University occupies. The city of New Orleans is a continuation of an indigenous trade hub on the Mississippi River, known for thousands of years as Bulbancha. Native peoples have lived on this land since time immemorial, and the resilient voices of Native Americans remain an inseparable part of our local culture. With gratitude and honor, we acknowledge the indigenous nations that have lived and continue to thrive here.

## Author Contributions

JML, AM, and KGF designed the research and wrote the paper. JML and AM performed the research and analyzed data. KGF funded the research and provided supervision.

## Data Accessibility and Benefit-Sharing

Phenotypic data and code is available on GitHub at the following repository: https://github.com/jill-love/Love-Mahesh-Ferris-2026

Genomic sequence data can be found in the NCBI BioSample database under accession number PRJNA1372734.

Benefits Generated: Benefits from this research accrue from the sharing of our data and results on public databases as described above.

