## supplemental for "The genomic basis of local adaptation to photoperiod across altitude in a self-fertilizing monkeyflower"

**Supplementary Figures**

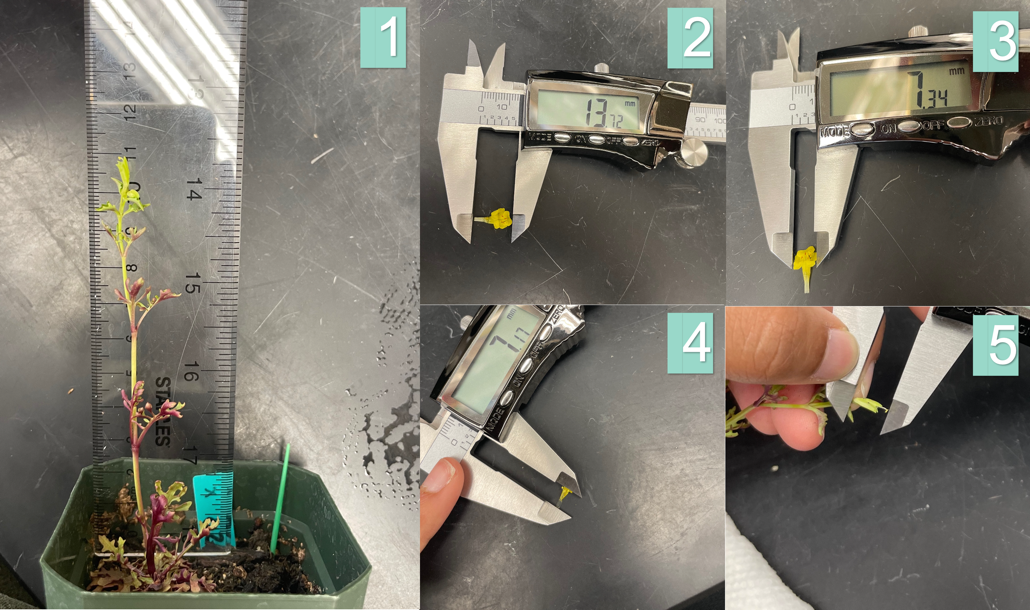

**Figure S1.** Phenotypic trait sampling in *M. laciniatus*.

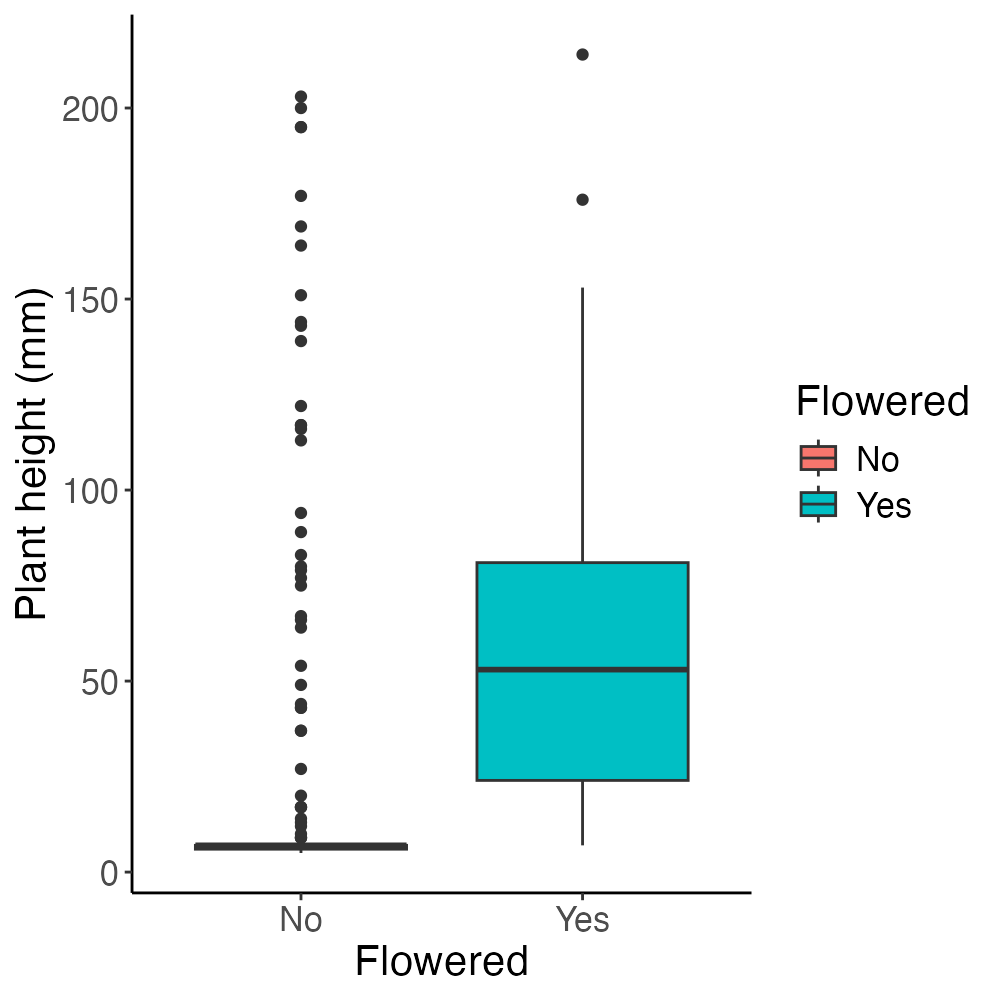

**Figure S2.** Plant height of flowering and non-flowering F_2_ individuals under the short-day 11 hour photoperiod.

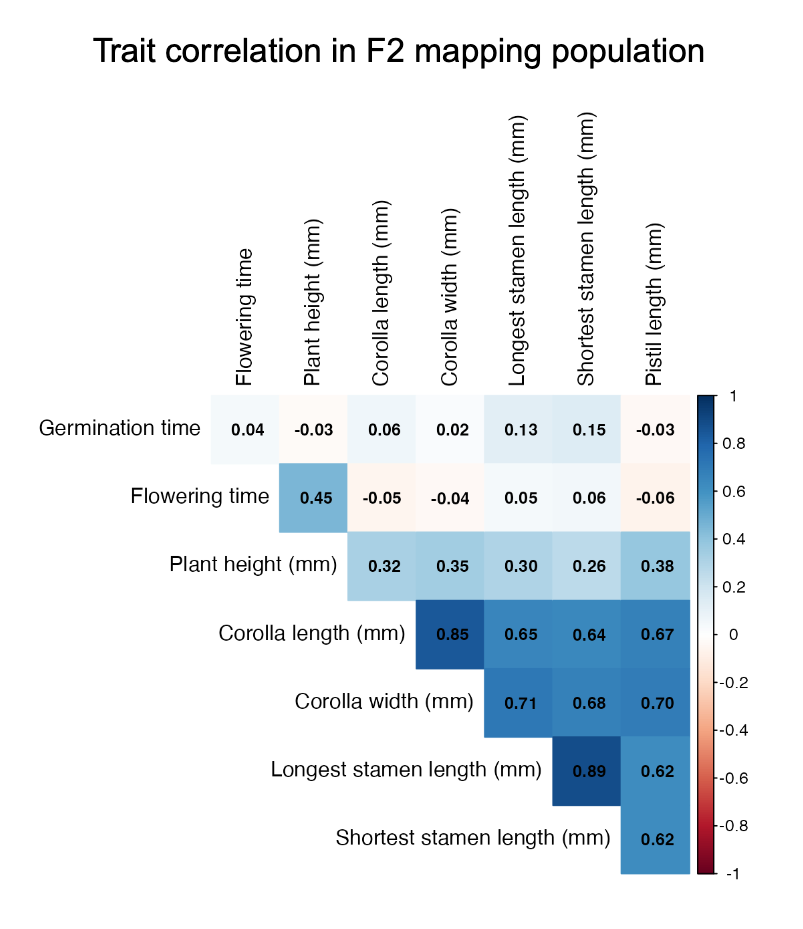

**Figure S3.** Correlation matrix of traits measured in the F_2_ mapping population. Values in squares represent Pearson correlation coefficients (*r*).

**Supplementary Tables**

Table S1. Expected and observed number of F_2_ individuals to flower in short-days if controlled by a single dominant mendelian locus.

|  | **Expected** | **Observed** |
| --- | --- | --- |
| **Absent** | 363 | 381 |
| **Present** | 121 | 103 |
| **Total** | 484 | 484 |

Table S2. Annotated list of candidate loci from 46 *G*-statistic clustered peaks. Gene functions derived from the TAIR database (Rhee et al., 2003).

| **Chr** | **Gene** | **bp_start** | **bp_end** | **Function (Arabidopsis homolog)** |
| --- | --- | --- | --- | --- |
| Chr_02 | MgTOL.B0865 | 6945009 | 6950051 | Encodes a protein kinase that phosphorylates histone H3 at Thr3 and Thr11 and plays a role in mitotic cell division. |
| Chr_02 | MgTOL.B0867 | 6929795 | 6932840 | NEAP2 is a member of a small family containing coiled-coil domains, a nuclear localization signal and a C-terminal predicted transmembrane domain. It localizes to the nuclear periphery. Mutants have altered nuclear morphology and chromatin structure. |
| Chr_02 | MgTOL.B0986 | 9285416 | 9290567 | RECQ helicase l1;(source:Araport11) |
| Chr_02 | MgTOL.B1053 | 10230256 | 10231459 | Encodes a putative RING-H2 zinc finger protein ATL6 (ATL6). |
| Chr_02 | MgTOL.B1053 | 10230256 | 10231459 | Encodes CNI1 (Carbon/Nitrogen Insensitive1) (also named as ATL31), a RING type ubiquitin ligase that functions in the Carbon/Nitrogen response for growth phase transition in Arabidopsis seedlings. |
| Chr_02 | MgTOL.B1054 | 10234498 | 10236197 | Protein kinase superfamily protein;(source:Araport11) |
| Chr_02 | MgTOL.B1055 | 10236664 | 10240691 | Phosphoglycerate mutase family protein;(source:Araport11) |
| Chr_02 | MgTOL.B1056 | 10245086 | 10249788 | Specifically deaminates (de)guanosine to produce xanthosine with high specificity, which is further converted to xanthine, a key intermediate in purine metabolism and nitrogen recycling. |
| Chr_02 | MgTOL.B1096 | 11374562 | 11374833 | encodes a chloroplast ribosomal protein L20, a constituent of the large subunit of the ribosomal complex |
| Chr_02 | MgTOL.B1277 | 14607182 | 14611425 | delta-latroinsectotoxin-Lt1a protein;(source:Araport11) |
| Chr_08 | MgTOL.H0795 | 5965942 | 5968280 | Ribosomal protein L17 family protein;(source:Araport11) |
| Chr_08 | MgTOL.H2014 | 17177812 | 17182215 | Encodes a homolog of animal DJ-1 superfamily protein. In the A. thaliana genome, three genes encoding close homologs of human DJ-1 were identified AT3G14990 (DJ1A), AT1G53280 (DJ1B) and AT4G34020 (DJ1C). Among the three homologs, DJ1C is essential for chloroplast development and viability. It exhibits glyoxalase activity towards glyoxal and methylglyoxal. |
| Chr_08 | MgTOL.H2045 | 17387823 | 17391573 | The Arabidopsis protein AtGGH1 is a gamma-glutamyl hydrolase cleaving pentaglutamates to yield di- and triglutamates. The enzyme is involved in the tetrahydrofolate metabolism and located to the vacuole. |
| Chr_08 | MgTOL.H2045 | 17387823 | 17391573 | gamma-glutamyl hydrolase 3;(source:Araport11) |
| Chr_08 | MgTOL.H2045 | 17387823 | 17391573 | The Arabidopsis protein AtGGH2 is a gamma-glutamyl hydrolase acting specifically on monoglutamates. The enzyme is involved in the tetrahydrofolate metabolism and located to the vacuole. |
| Chr_08 | MgTOL.H2135 | 18685798 | 18690570 | RNA-binding protein, putative, similar to RNA-binding protein GB:AAA86641 GI:1174153 from (Arabidopsis thaliana).Contains PAB2 domain which facilitates binding to PABC proteins. |
| Chr_08 | MgTOL.H2142 | 18770062 | 18772274 | Encodes a ROP/RAC effector, designated interactor of constitutive active ROPs 1 (ICR1), that interacts with GTP-bound ROPs. ICR1 is a scaffold mediating formation of protein complexes that are required for cell polarity. ICR1 is comprised of coiled-coil domains and forms complexes with itself and the exocyst vesicle-tethering complex subunit SEC3. |
| Chr_08 | MgTOL.H2142 | 18770062 | 18772274 | Encodes RIP2 (ROP interactive partner 2), a putative Rho protein effector, interacting specifically with the active form of ROPs (Rho proteins of plants). |
| Chr_08 | MgTOL.H2143 | 18773581 | 18777373 | Encodes a gibberellin 2-oxidase that acts on C-19 gibberellins to deactivate them. AtGA2OX2 expression is responsive to cytokinin and KNOX activities. |
| Chr_08 | MgTOL.H2223 | 20294787 | 20298879 | S-adenosyl-L-methionine-dependent methyltransferases superfamily protein;(source:Araport11) |
| Chr_08 | MgTOL.H2418 | 23122445 | 23128717 | signal transducer, putative (DUF3550/UPF0682);(source:Araport11) |
| Chr_08 | MgTOL.H2419 | 23133697 | 23135566 | Histone variant H3. Associated with gene body methylation. |
| Chr_08 | MgTOL.H2420 | 23136369 | 23139704 | Ubiquitin carboxyl-terminal hydrolase family protein;(source:Araport11) |
| Chr_08 | MgTOL.H2421 | 23141072 | 23145169 | RNA-binding splicing factor; part of SWAP1-SFPS-RRC1 splicing factor complex modulates pre-mRNA splicing to promote photomorphogenesis. |
| Chr_08 | MgTOL.H2575 | 24179287 | 24185853 | Member of the microrchidia protein family which have been described as epigenetic regulators and plant immune mediators, contains a hallmark GHKL-type ATPase domain in N-terminus. |
| Chr_08 | MgTOL.H2575 | 24179287 | 24185853 | R-protein-interacting protein that localizes to endosomes and functions in resistance gene?mediated immunity. Belongs to the conserved Microrchidia (MORC) adenosine triphosphatase (ATPase) family, predicted to catalyze alterations in chromosome superstructure. Required for heterochromatin condensation and gene silencing.The expression of MORC1 requires the activity of Plant Mobile Domain proteins MAIN and MAIL1. |
| Chr_08 | MgTOL.H2576 | 24190165 | 24192207 | Encodes a protein of 231 amino acids with 51% identity to RTE1 over 209 amino acids. Interacts with RTE1 in planta and appears to function in same pathway to positively regulate ethylene signaling. |
| Chr_08 | MgTOL.H2578 | 24193465 | 24196432 | phosphatidylinositolglycan-like protein;(source:Araport11) |
| Chr_08 | MgTOL.H2579 | 24197432 | 24198806 | encodes a cytosolic thioredoxin that reduces disulfide bridges of target proteins by the reversible formation of a disulfide bridge between two neighboring Cys residues present in the active site. Thioredoxins have been found to regulate a variety of biological reactions in prokaryotic and eukaryotic cells. |
| Chr_11 | MgTOL.K1365 | 15814696 | 15817055 | Target gene of MIR2111-5p. The miR2111-TML/HOLT regulon is involved in control of lateral root initiation. |
| Chr_11 | MgTOL.K1365 | 15814696 | 15817055 | Galactose oxidase/kelch repeat superfamily protein;(source:Araport11) |
| Chr_11 | MgTOL.K1367 | 15819874 | 15821506 | ENY2 is a component of the deubiquitination module of the SAGA complex and is involved in the deubiquitination of histone 2B . |
| Chr_11 | MgTOL.K1368 | 15830552 | 15833289 | Component of the TOM complex involved in transport of nuclear-encoded mitochondrial proteins |
| Chr_13 | MgTOL.M1557 | 16233429 | 16236003 | S-adenosyl-L-methionine-dependent methyltransferases superfamily protein;(source:Araport11) |
| Chr_13 | MgTOL.M1558 | 16239961 | 16243163 | Encodes for a protein with ent-kaurene synthase B activity which catalyzes the second step in the cyclization of GGPP to ent-kaurene in the gibberellins biosynthetic pathway. |
| Chr_13 | MgTOL.M1560 | 16247944 | 16250103 | Aldolase-type TIM barrel family protein;(source:Araport11) |
| Chr_13 | MgTOL.M1561 | 16250898 | 16252168 | Carbohydrate-binding X8 domain superfamily protein;(source:Araport11) |
| Chr_13 | MgTOL.M1562 | 16253431 | 16255646 | Pentatricopeptide repeat (PPR) superfamily protein;(source:Araport11) |
| Chr_14 | MgTOL.N1986 | 16988670 | 16992655 | Eukaryotic aspartyl protease family protein;(source:Araport11) |
